# Behavioral and neural evidence for the underestimated attractiveness of faces synthesized using an artificial neural network

**DOI:** 10.1101/2023.02.07.527403

**Authors:** Satoshi Nishida

## Abstract

Despite recent advantages in artificial intelligence (AI), the potential human aversion to AI has not been dispelled yet. If such aversion degrades the human preference to AI-synthesized visual information, the preference should be reduced solely by the human belief that the information is synthesized by AI, independently of its appearance. To test this hypothesis, this study designed a task paradigm in which naïve participants rated the attractiveness of various faces synthesized using an artificial neural network, under the fake instruction that half of the faces were synthetic and the other half were real. This design allowed evaluating the effect of participants’ beliefs on their attractiveness ratings separately from the effect of facial appearance. In addition, to investigate the neural substrates of the belief effect, brain responses to faces were collected using fMRI during this task. It is found that participants’ ratings declined when the faces were believed to be synthetic. Furthermore, the belief changed the responsiveness of fMRI signals to facial attractiveness in the right fusiform cortex. These behavioral and neural findings support the notion that the human preference to visual information becomes lower solely due to the beliefs that the information is synthesized by AI.

## 1. Introduction

Recent advances in artificial intelligence (AI) technology have been facilitated by the advent of artificial neural networks using deep learning (Goodfellow et al., 2016; LeCun et al., 2015). One of its most prominent advances is image synthesis, whereby an infinite number of fake images (e.g., faces) are synthesized by artificial neural networks such as the generative adversarial networks (Goodfellow et al., 2014; Karras, Laine, & Aila, 2019). These synthetic images are realistic enough to make it difficult for humans to discriminate them from real images (Nightingale & Farid, 2022).

Does such AI-synthesized visual information attract humans as much as real images? Previous AI/robot research has emphasized the effect of AI/robot appearance on its attractiveness for humans (Abubshait & Wiese, 2017; DiSalvo et al., 2002; Gong, 2008; Kanda et al., 2008; Koda & Maes, 1996; McDonnell et al., 2012). For instance, the “uncanny valley” hypothesis, which explains a well-known cognitive phenomenon of human negative affection against artificial objects (Mori et al., 2012), indicates that the attractiveness of artificial objects abruptly drops as the object appearance approaches that of humans. In addition, they predict that an object’s attractiveness improves when their appearance becomes so similar to the real object that humans cannot discriminate between them. If this prediction were true, the human subjective attractiveness of AI-synthesized visual information would be determined solely by the appearance of the visual information.

Despite recent advances in AI technology, there is still aversion to AI (e.g., AI will get out of control; AI will dominate humankind) (Li & Huang, 2020); this aversion may stem from the representation of AI in fictional stories, such as sci-fi books and movies [c.f. Frankenstein complex; (Asimov, 1964)]. Such aversion is one of the most crucial factors decreasing human trust in AI (Siau & Wang, 2018). Therefore, such pre-existing aversion may degrade the subjective attractiveness of AI-synthesized visual information. If this is the case, the attractiveness should be reduced solely by the human belief that the information is synthesized by AI, independently of its appearance.

To investigate this hypothesis, I designed a cognitive task to examine whether the subjective attractiveness of AI-synthesized visual images is affected by the participants’ beliefs that the images are synthetic, even after eliminating the effects of image appearance (Fig.1). In this attractiveness-rating task, human participants rated the attractiveness of various face images under two instruction conditions (they were told that the images were either real or synthetic) even though all images were synthesized by an artificial neural network. The behavioral effect of the belief was evaluated by the changes in attractiveness ratings obtained across conditions.

**Fig. 1.**
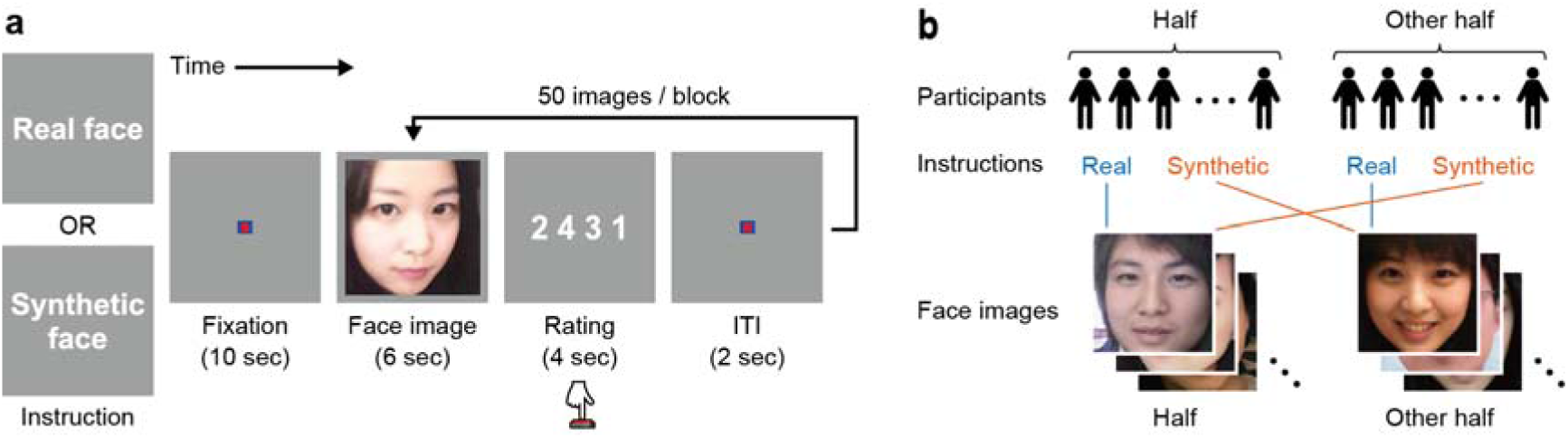
Attractiveness-rating task. (a) Task procedure. The task consisted of six blocks for which the subjective attractiveness of 50 face images was manually rated. At the beginning of each block, information on whether the face images presented in that block were real (real-face condition) or synthetic (synthetic-face condition) was provided to the participant. After a 10 s fixation period, an initial face image was shown for 6 s. Then, four digits (“1,” “2,” “3,” and “4”) were presented in a random order for a fixed period of 4 s to prompt the participant to provide their attractiveness-rating score for the face shown by pressing the corresponding button. After the digits disappeared, a 2 s inter-trial interval (ITI) was imposed before the next image presentation. Each participant completed three blocks under the real-face condition and another three blocks under the synthetic-face condition in an interleaved manner. The face image shown in this figure is AI-synthesized and not of real people. (b) Face condition associations were counterbalanced across participants. All 300 face images were divided into two different sets of 150 images (sets 1 and 2). Half of the participants performed attractiveness rating on set 1 believing them to be real faces and on set 2 to be synthetic faces whereas the other half of participants did the opposite. This design enabled to collect attractiveness-rating scores on the same images under both the real- and synthetic conditions at the group level. The face images shown in this figure are AI-synthesized and not of real people.

In addition, I also explored the neural correlates of these changes. To this end, brain responses to face images during the attractiveness-rating task were measured using fMRI. After localizing the brain regions indicating the perception of facial attractiveness, I examined the instruction-dependent changes in facial-attractiveness signals in these regions from the following two perspectives: (1) which regions change their signals depending on the instructions and (2) which show instruction-dependent changes according to the individual variation in attractiveness ratings?

## 2. Materials and Methods

### 2.1. Participants

Two separate groups of 30 (15 females; age [mean ± SD] = 25.0 ± 8.1SD years) and 60 (30 females; age [mean ± SD] = 23.3 ± 5.1SD years) healthy Japanese participants were recruited for the preliminary survey and fMRI experiments, respectively. All had normal or corrected-to-normal vision. Written informed consent was obtained from all participants. The experimental protocol was approved by the ethics and safety committees of National Institute of Information and Communications Technology.

### 2.2. Stimuli

Three hundred different color face images were synthesized using StyleGAN2 (Karras, Laine, Aittala, et al., 2019), an upgraded version of StyleGAN (Karras, Laine, & Aila, 2019). A version of StyleGAN2 implementation available online (https://github.com/lucidrains/stylegan2-pytorch) was used with default parameters except for the output image size (256 × 256 pixels). A StyleGAN2 network was trained using the Asian Face Age Dataset [AFAD; (Niu et al., 2016)] since the participants were more familiar with Asian faces than faces from other ethnicities. After the training, 300 face images (150 females and 150 males) were manually selected from a large set of images synthesized by the trained network. The selection was performed such that all selected images appeared like real faces, that is, they did not contain anything unnatural element (e.g., distortion, discontinuity, unnatural color, etc.). This arbitrariness of the selection criterion did not affect ratings, because the attractiveness ratings were collected from the same 300 images being considered both real and synthetic and compared (see below).

### 2.3. Preliminary survey

Attractiveness was rated by each participant on two separate image sets including the same number of faces (i.e., 150 each) under different instructions, and the mean rating between image sets was compared. Since a large difference of original attractiveness (i.e., without any instructions) between image sets would complicate the rating comparison within each participant, the image sets were split in half according to a preliminary survey.

In the survey, each participant rated the attractiveness of 300 faces presented sequentially on a computer monitor at a distance of 60 cm from the participant. Each face image was presented at the center of the monitor (15.9 × 15.9 degree of visual angle [dva]) with a horizontal 7-scale rating bar shown underneath the image. The participant selected among 1 to 7 to provide the attractiveness score by a mouse click and then pressed a selection button. The image remained until the participant’s button press and changed to the next image immediately thereafter.

The attractiveness ratings collected for each image were averaged across participants. According to this average attractiveness rating, all 300 faces were divided into two sets. First, the distribution of the average rating values was grouped into 15 equally spaced bins. In each bin, half of each of the corresponding male and female faces was randomly drawn from all faces in that bin. This procedure yielded two sets of 75 male and 75 female faces with an almost equal distribution of attractiveness ratings.

### 2.4. Attractiveness-rating task

Each participant performed an attractiveness-rating task inside an MRI scanner. In this task, each participant was asked to rate the attractiveness of the 300 faces presented separately in each of six blocks (i.e., 50 faces each). Although the set of 50 faces in each block was identical for all participants, the order of the faces presented in each block was randomized across participants.

At the beginning of each block, an instruction at the center of the screen was provided to inform that all the faces presented in that block were either real (real-face condition) or synthetic (synthetic-face condition). After commencing fMRI scanning, a 10 s fixation period was imposed to avoid the undesirable effects of hemodynamic signal instability. The fMRI data collected during this fixation period was discarded from the analysis. Then, a face image (7.68 × 7.68 dva) was presented at the center of the screen for 6 s; participants’ eye movements were not restricted. After the face image disappeared, four digits (“1,” “2,” “3,” and “4”) equally spaced horizontally were presented at the center of the screen for 4 s. The digit order was randomized from trial to trial to avoid the fixed association of each digit with a specific finger. Participants were asked to choose one of the four digits as the attractiveness score of the face and press a button in the same position as that of the chosen digit in the screen. Participants were allowed to press a button multiple times within the 4 s period. The last pressed button was regarded as the participants’ choice. A button press beyond this period was not accepted. After the rating period, a 2 s fixation period was imposed as an inter-trial interval before face presentation in the next trial.

Each participant completed six interleaved blocks, three under the real-face condition and three under the synthetic-face condition. The block order under these conditions was counterbalanced across participants. Half of the participants performed blocks 1, 3, 5 under the real-face condition and blocks 2, 4, 6 under the synthetic-face condition whereas the other half did the opposite. This design enabled to obtain the attractiveness rating of all faces under both conditions at the group level.

### 2.5. Behavioral analysis

The effect of the different instructions on facial attractiveness ratings was evaluated at the level of individual faces and individual participants. For the face-level analysis of the instruction effect, the attractiveness ratings for each face were averaged over participants separately for the synthetic- and real-face conditions. Then, the condition difference in the average ratings for individual faces was calculated by subtracting the ratings in the synthetic-face condition from those in the real-face condition. Since positive values of the rating difference reflected the participants’ underestimation of facial attractiveness in the synthetic-face condition, this rating difference was called *underestimation score*. In the participant-level analysis of instruction effects, the attractiveness ratings in a given participant were averaged over faces separately for the synthetic- and real-face conditions. Then, the underestimation scores were calculated by subtracting the ratings obtained in the synthetic-face condition from those in the real-face condition. These face- and participant-wise underestimation scores were used to evaluate instruction effects separately at the face- and participant levels, respectively.

### 2.6. MRI data collection

Functional and anatomical MRI data were collected using a 3T Siemens MAGNETOM Vida scanner (Siemens, Germany) with a 64-channel Siemens volume coil. Functional data were collected during the attractiveness-rating task using a multiband gradient echo EPI sequence ((Moeller et al., 2010); repetition time [TR] = 2,000 ms; echo time [TE] = 30 ms; flip angle = 75°; voxel size = 2 × 2 × 2 mm; matrix size = 100 × 100; number of slices = 75; multiband factor = 3). Anatomical data were collected using a T1-weighted MPRAGE sequence (TR = 2,530 ms; TE = 3.26 ms; flip angle = 9°; voxel size = 1 × 1 × 1 mm; matrix size = 256 × 256; number of slices = 208).

### 2.7. MRI data preprocessing

Results included in this manuscript come from preprocessing performed using fMRIPrep 20.2.6 ((Esteban et al., 2018, 2019); RRID:SCR_016216), which is based on Nipype 1.7.0 ((K. Gorgolewski et al., 2011; K. J. Gorgolewski et al., 2018); RRID:SCR_002502).

#### Anatomical data preprocessing

A total of 1 T1-weighted (T1w) images were found within the input BIDS dataset. The T1-weighted (T1w) image was corrected for intensity non-uniformity (INU) with N4BiasFieldCorrection (Tustison et al., 2010), distributed with ANTs 2.3.3 ((Avants et al., 2008); RRID:SCR_004757), and used as T1w-reference throughout the workflow. The T1w-reference was then skull-stripped with a Nipype implementation of the antsBrainExtraction.sh workflow (from ANTs), using OASIS30ANTs as target template. Brain tissue segmentation of cerebrospinal fluid (CSF), white-matter (WM) and gray-matter (GM) was performed on the brain-extracted T1w using fast (FSL 5.0.9, RRID:SCR_002823, (Zhang et al., 2001)). Brain surfaces were reconstructed using recon-all (FreeSurfer 6.0.1, RRID:SCR_001847, (Dale et al., 1999)), and the brain mask estimated previously was refined with a custom variation of the method to reconcile ANTs-derived and FreeSurfer-derived segmentations of the cortical gray-matter of Mindboggle (RRID:SCR_002438, (Klein et al., 2017)). Volume-based spatial normalization to one standard space (MNI152NLin2009cAsym) was performed through nonlinear registration with antsRegistration (ANTs 2.3.3), using brain-extracted versions of both T1w reference and the T1w template. The following template was selected for spatial normalization: ICBM 152 Nonlinear Asymmetrical template version 2009c [(Fonov et al., 2009), RRID:SCR_008796; TemplateFlow ID: MNI152NLin2009cAsym],

#### Functional data preprocessing

For each of the 6 BOLD runs found per subject (across all tasks and sessions), the following preprocessing was performed. First, a reference volume and its skull-stripped version were generated using a custom methodology of fMRIPrep. A B0-nonuniformity map (or fieldmap) was estimated based on a phase-difference map calculated with a dual-echo GRE (gradient-recall echo) sequence, processed with a custom workflow of SDCFlows inspired by the epidewarp.fsl script and further improvements in HCP Pipelines (Glasser et al., 2013). The fieldmap was then co-registered to the target EPI (echo-planar imaging) reference run and converted to a displacements field map (amenable to registration tools such as ANTs) with FSL’s fugue and other SDCflows tools. Based on the estimated susceptibility distortion, a corrected EPI (echo-planar imaging) reference was calculated for a more accurate co-registration with the anatomical reference. The BOLD reference was then co-registered to the T1w reference using bbregister (FreeSurfer) which implements boundary-based registration (Greve & Fischl, 2009).

Co-registration was configured with six degrees of freedom. Head-motion parameters with respect to the BOLD reference (transformation matrices, and six corresponding rotation and translation parameters) are estimated before any spatiotemporal filtering using mcflirt (FSL 5.0.9, (Jenkinson et al., 2002)). BOLD runs were slice-time corrected to 0.95s (0.5 of slice acquisition range 0s-1.9s) using 3dTshift from AFNI 20160207 ((Cox & Hyde, 1997), RRID:SCR_005927). The BOLD time-series (including slice-timing correction when applied) were resampled onto their original, native space by applying a single, composite transform to correct for head-motion and susceptibility distortions. These resampled BOLD time-series will be referred to as preprocessed BOLD in original space, or just preprocessed BOLD. The BOLD time-series were resampled into standard space, generating a preprocessed BOLD run in MNI152NLin2009cAsym space. First, a reference volume and its skull-stripped version were generated using a custom methodology of fMRIPrep. Several confounding time-series were calculated based on the preprocessed BOLD: framewise displacement (FD), DVARS and three region-wise global signals. FD was computed using two formulations following Power (absolute sum of relative motions, (Power et al., 2014)) and Jenkinson (relative root mean square displacement between affines, (Jenkinson et al., 2002)). FD and DVARS are calculated for each functional run, both using their implementations in Nipype (following the definitions by (Power et al., 2014)). The three global signals are extracted within the CSF, the WM, and the whole-brain masks. Additionally, a set of physiological regressors were extracted to allow for component-based noise correction (CompCor, (Behzadi et al., 2007)).

Principal components are estimated after high-pass filtering the preprocessed BOLD time-series (using a discrete cosine filter with 128s cut-off) for the two CompCor variants: temporal (tCompCor) and anatomical (aCompCor). tCompCor components are then calculated from the top 2% variable voxels within the brain mask. For aCompCor, three probabilistic masks (CSF, WM and combined CSF+WM) are generated in anatomical space. The implementation differs from that of Behzadi et al. in that instead of eroding the masks by 2 pixels on BOLD space, the aCompCor masks are subtracted a mask of pixels that likely contain a volume fraction of GM. This mask is obtained by dilating a GM mask extracted from the FreeSurfer’s aseg segmentation, and it ensures components are not extracted from voxels containing a minimal fraction of GM. Finally, these masks are resampled into BOLD space and binarized by thresholding at 0.99 (as in the original implementation). Components are also calculated separately within the WM and CSF masks. For each CompCor decomposition, the k components with the largest singular values are retained, such that the retained components’ time series are sufficient to explain 50 percent of variance across the nuisance mask (CSF, WM, combined, or temporal). The remaining components are dropped from consideration. The head-motion estimates calculated in the correction step were also placed within the corresponding confounds file. The confound time series derived from head motion estimates and global signals were expanded with the inclusion of temporal derivatives and quadratic terms for each (Satterthwaite et al., 2013). Frames that exceeded a threshold of 0.5 mm FD or 1.5 standardised DVARS were annotated as motion outliers. All resamplings can be performed with a single interpolation step by composing all the pertinent transformations (i.e. head-motion transform matrices, susceptibility distortion correction when available, and co-registrations to anatomical and output spaces). Gridded (volumetric) resamplings were performed using antsApplyTransforms (ANTs), configured with Lanczos interpolation to minimize the smoothing effects of other kernels (Lanczos, 1964). Non-gridded (surface) resamplings were performed using mri_vol2surf (FreeSurfer).

Many internal operations of fMRIPrep use Nilearn 0.6.2 ((Abraham et al., 2014), RRID:SCR_001362), mostly within the functional processing workflow. For more details of the pipeline, see the section corresponding to workflows in fMRIPrep’s documentation (https://fmriprep.readthedocs.io/en/latest/workflows.html).

The above text was automatically generated by fMRIPrep with the express intention that users should copy and paste this text into their manuscripts unchanged. It is released under the CC0 license.

### 2.8. General linear model (GLM) analysis

Further data analysis was performed using Nilearn (Abraham et al., 2014). Prior to the GLM analysis of the preprocessed fMRI data in MNI152NLin2009cAsym space, additional preprocessing, including linear trend removal, high-pass filtering (128 Hz), and z-scoring, was performed within each run. The trials in which participants pressed no button within the 4 s rating period were discarded from the analysis.

In first-level analysis, GLM with an event-related design was applied to the preprocessed fMRI data. Prior to model fitting, fMRI signals were spatially smoothed using a Gaussian kernel with a full-width at half maximum of 10 mm. The fMRI signals during the 6 s period of face presentation in each trial were modeled as boxcars and convolved with a canonical hemodynamic response function (Friston et al., 1998). First-level GLM included two regressors of interest (synthetic- and real-face conditions) using each participant’s attractiveness rating on each face to model the linear parametric modulations of task instruction according to individual ratings. The model contained nuisance regressors of six head-motion parameters, framewise displacement, and the first six aCompCor components. The obtained β-coefficient estimates for each regressor reflect voxel responsiveness to facial attractiveness in the synthetic- or real-face conditions.

These two β-coefficient estimates were entered into second-level, random-effects analysis accounting for the between-participant variance. To localize brain regions encoding facial attractiveness regardless of task conditions, a one-sample t-test was performed in each voxel using the mean of the two β-coefficient estimates for each participant. The resulting t values were subsequently transformed into z scores to generate a statistical parametric map for the effect of facial attractiveness on each voxel response. Then, cluster-level inference for family-wise error (FWE) control was performed using threshold free cluster enhancement (TFCE) analysis (Smith & Nichols, 2009), which allows cluster-level FWE correction without explicit use of an a priori cluster-forming p-value threshold. For controlling FWE rate, a p < 0.05 after TFCE cluster-level correction was used as threshold of statistical significance for the statistical parametric map from the second-level analysis.

### 2.9. Region of interest (ROI) analysis

For further acquisition of neural evidence for the underestimated attractiveness of synthetic faces, ROIs corresponding to voxel clusters signaling facial attractiveness were determined according to the second-level GLM analysis. Among voxels with significant t values in the second-level analysis (p < 0.05, cluster-level FWE corrected), separate regions with ≥50 connected voxels (i.e., 400 mm^3^) were extracted using the function *connected_regions* in Nilearn and regarded as ROIs. To identify the brain regions involved in the attractiveness underestimation for synthetic faces, two types of ROI-based analysis were performed. The first, a simpler analysis, aimed to explore brain regions changing their group-average responsiveness to faces under the different instructions. To this end, regional responsiveness to faces under each task condition was estimated in each ROI for each participant. For this estimation, the β-coefficient estimates for the regressor of each task condition from the first-level GLM analysis were used after averaging over all voxels in a given ROI. Then, in each ROI, the regional responsiveness under the two task conditions was compared using a paired t-test with Bonferroni correction for multiple comparisons.

The second analysis aimed to examine the behavioral correlates of the attractiveness underestimation for synthetic faces in each brain region. If a given brain region was involved in the attractiveness underestimation, the neural signal change between task conditions in that region should co-vary with the attractiveness-rating change across conditions (i.e., underestimation score). For this analysis, the regional responsiveness obtained for each task-condition was evaluated by the voxel-average β-coefficient estimates for each participans from the first-level GLM analysis. Then, the regional responsiveness for the synthetic-face condition was subtracted from that for the real-face condition to estimate the neural signal change between conditions. Then, in each ROI, the co-variation between the responsiveness changes and underestimation scores was examined by a Pearson correlation across participants with Bonferroni correction for multiple comparisons.

## 3. Results

### 3.1. Underestimated facial attractiveness under the instruction that faces are synthetic

All 60 participants rated the attractiveness of 300 face images between 1–4 through a button press during the attractiveness-rating task performed in an MRI scanner (Fig. 1). One hundred and fifty images were instructed to be synthetic (synthetic-face condition) whereas the other 150 were instructed to be real (real-face condition). In a preliminary survey, the two sets of face images to be rated under different task conditions had been divided such that the images in each set had comparable average attractiveness ratings under no instructions (see Materials and Methods). In fact, the attractiveness ratings in each condition of the attractiveness-rating task did not significantly differ between image sets (mean ± standard error of mean [SEM]: synthetic-face condition, 2.26 ± 0.06 and 2.19 ± 0.06, two-sample t-test, p = 0.42; real-face condition, 2.21 ± 0.05 and 2.18 ± 0.06, p = 0.86).

The participants’ underestimation of facial attractiveness in the synthetic-face condition relative to the real-face condition was evaluated using underestimation scores (see Materials and Methods). The face-wise underestimation scores, which assessed the face-level effect of instruction, were significantly higher than zero over all faces (Fig. 2a; one-sample t-test, p < 0.001; Wilcoxon sign-rank test, p < 0.005). This face-level analysis also provided information about the faces resulting in higher underestimated attractiveness in the synthetic-face condition (Fig. S1a). Despite the large variability in underestimation scores across faces (Fig. 2a), no clear difference in facial attributes was observed based on facial attractiveness underestimation. Male and female faces resulted in no significant differences in underestimation scores (Fig. S1b). Further, the face-wise underestimation scores were unlikely to co-vary with the attractiveness of the faces itself (Fig. S1c).

**Fig. 2.**
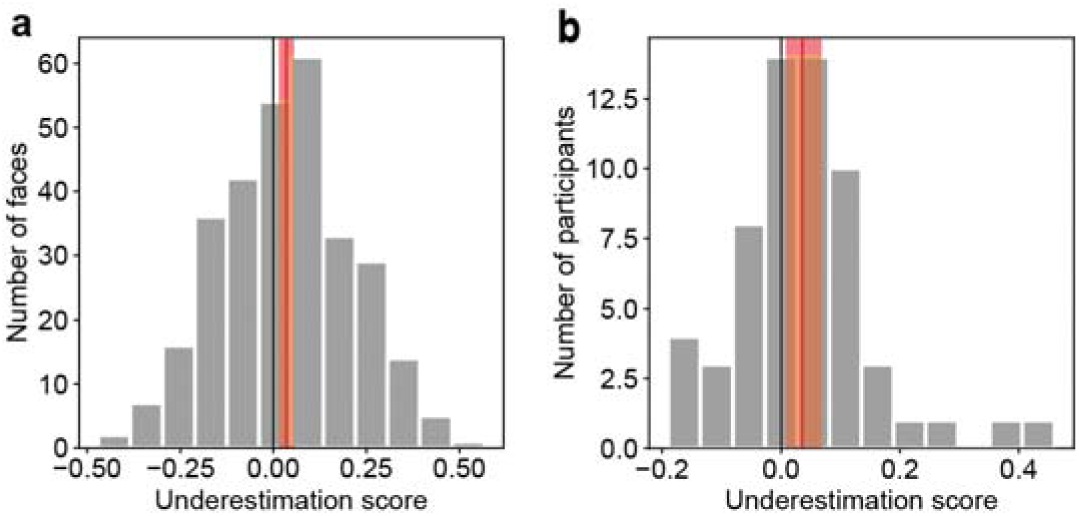
Underestimated attractiveness ratings for synthetic faces. The underestimation of attractiveness for synthetic faces relative to real ones was evaluated by subtracting the attractiveness ratings in the synthetic-face condition from those in the real-face condition (*underestimation score*). The underestimation score was computed per face (a) and per participant (b). Red vertical lines denote the average underestimation score. The red area around each line indicates the 95% confidence interval (CI).

In addition, the participant-wise underestimation scores, which assessed the participant-level effect of instruction, were also significantly higher than zero over all 60 participants (Fig. 2b one-sample t-test, p < 0.05; Wilcoxon signed rank test, p < 0.05). Taken together, these results indicate that facial attractiveness was underestimated in the synthetic-face condition.

Facial attractiveness is affected by other personal factors, such as age, sex, and sexual orientation (Foos & Clark, 2011; Hahn et al., 2016; Mitrovic et al., 2016). However, the participant-wise attractiveness underestimation of synthetic faces cannot be explained by these factors. There was no significant correlation between underestimation scores and age (Pearson correlation, r = 0.029, p = 0.83), between sexes, or different sexual orientations (two-sample t-test, p = 0.55 and 0.57, respectively). Taken together, these results provide behavioral evidence supporting that facial attractiveness is underestimated solely due to the human belief that the faces are synthetic.

### 3.2. Neural signals associated with attractiveness underestimation

To identify the neural substrates involved in the attractiveness underestimation of synthetic faces, I explored brain regions showing neural signal changes are associated with changes in attractiveness rating. For this purpose, neural signals were collected with fMRI during the attractiveness-rating task and analyzed using GLM to estimate single-voxel responsiveness to facial attractiveness under synthetic- and real-face conditions. Then, I aimed to localize the brain regions responding to facial attractiveness and whose responsiveness depends on underestimation scores (for more details, see Materials and Methods).

First, I identified the brain regions producing signals associated with facial attractiveness regardless of task conditions. To this end, the estimated responsiveness to facial attractiveness in each voxel was captured across conditions and statistically mapped in the brain through group-level random-effect analysis (Fig. 3). Multiple regions showed significantly positive responsiveness to facial attractiveness (p < 0.05, cluster-level FWE corrected), including the bilateral fusiform cortex, the right parieto-occipital cortex, the right intraparietal cortex, the right ventrolateral prefrontal cortex, the right insular cortex, the left orbitofrontal cortex, and the cerebellum (Fig. S2); in contrast, there was no voxel showing significantly negative responsiveness (p > 0.05, cluster-level FWE corrected). These regions consistently overlapped with previously reported brain regions considered as neural substrates of facial attractiveness judgments on real faces (Aharon et al., 2001; Bzdok et al., 2011; Iaria et al., 2008; Ishai, 2007; Mende-Siedlecki et al., 2013; O’Doherty et al., 2003; Winston et al., 2007). These eight regions were considered as ROIs for further analysis.

**Fig. 3.**
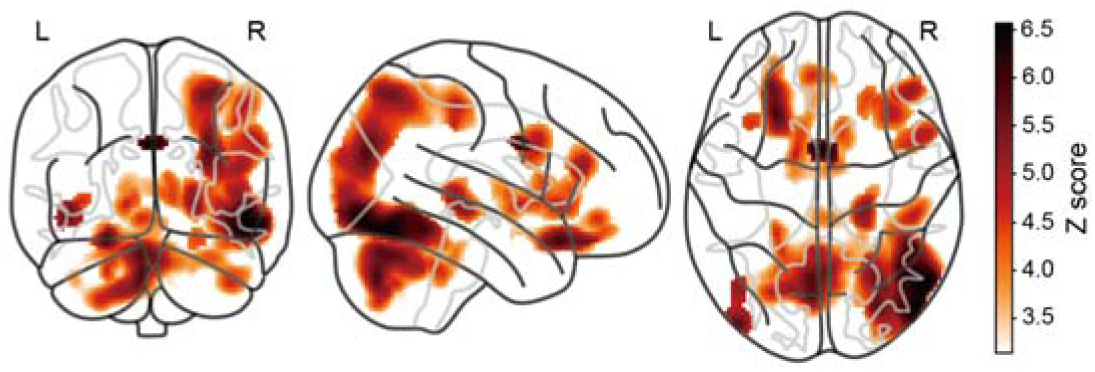
Brain regions encoding facial attractiveness. The voxel-wise statistics (z scores) of group-level analysis for the effect of facial attractiveness on voxel responses are mapped in a glass brain. Darker colors indicate voxels showing strong responsiveness to facial attractiveness. Only voxels with significant responsiveness are displayed (p < 0.05, cluster-level FWE corrected). No voxel showed negative responsiveness to facial attractiveness. L, left hemisphere; R, right hemisphere.

Next, I examined condition-dependent responsiveness differences associated with the behavioral changes in attractiveness ratings in each of the eight ROIs, from the following two perspectives. First, I explored brain regions showing changes in group-average responsiveness to facial attractiveness according to task conditions. To this end, the estimated responsiveness to facial attractiveness was compared, at the group level, between synthetic and real conditions after averaging over all voxels within each ROI. However, no ROI showed significant differences in estimated responsiveness between conditions (Fig. S3; paired t-test, P > 0.36, uncorrected).

Although the obtained behavioral results revealed a significant condition-dependent difference in attractiveness ratings, this effect was small at the group level (Fig. 2). Thus, this small behavioral difference might not be clearly reflected in group-average responsiveness changes estimated in these brain regions. Given the large behavioral differences across participants (Fig. 2b), the individual variability may be a more suitable target for exploring the neural correlates of attractiveness underestimation.

Therefore, I sought to identify the brain regions associated with this individual variability in attractiveness underestimation. To this end, I calculated condition-dependent differences in the estimated responsiveness to facial attractiveness within each ROI for each participant and tested a correlation between the estimated-responsiveness differences and the underestimation scores across participants. After correcting for multiple comparisons, this neural-behavior correlation was significant only in the right fusiform cortex (Fig. 4; Pearson r = 0.45, p < 0.005, Bonferroni corrected). This result indicates that among the brain regions signaling facial attractiveness, the signal change only in the right fusiform cortex is associated with the rating change between task conditions, providing neural evidence underlying attractiveness underestimation due to the belief that faces are synthetic.

**Fig. 4.**
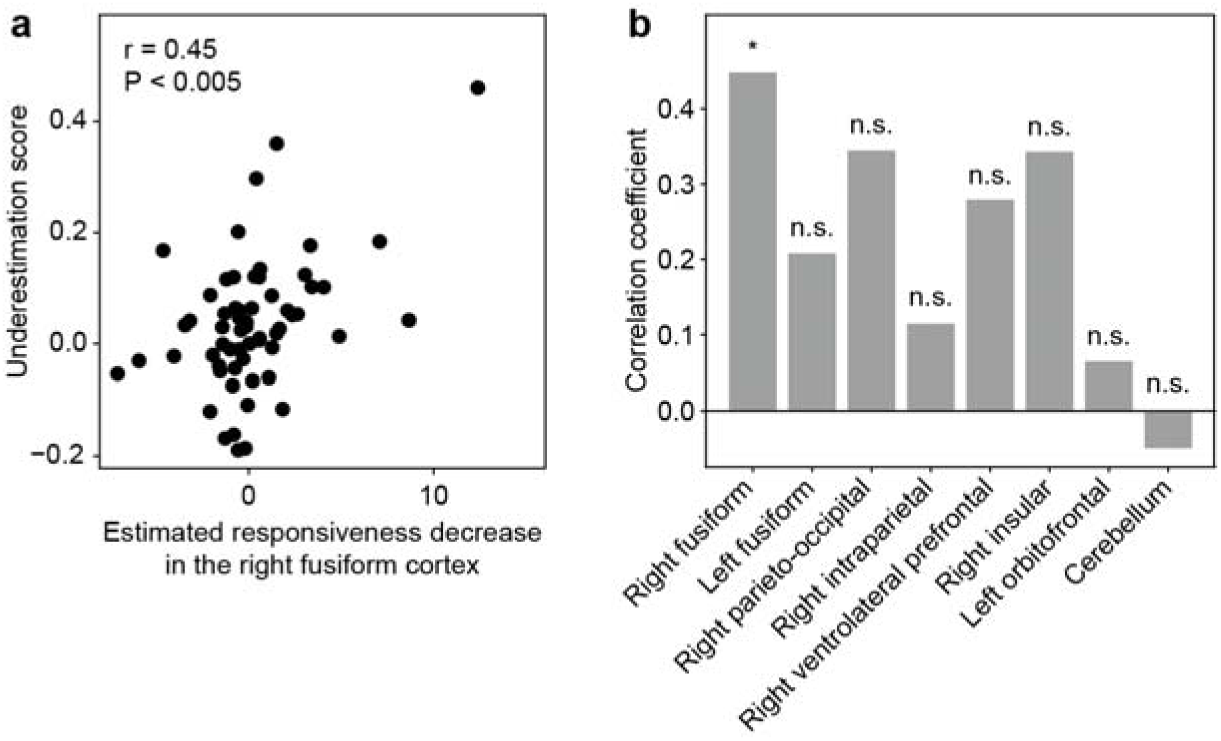
Association between the individual variability in neural responsiveness decrease and that in attractiveness underestimation. (a) Correlation between underestimation scores and estimated responsiveness decrease in the right fusiform cortex. For each participant, I calculated a decrease in estimated neural responsiveness to facial attractiveness as well as an underestimation score. Then, the correlation of these two measures was evaluated across participants. Each dot represents a single participant. (b) Summary of neural-behavior correlation in each of the eight ROIs. The Pearson correlation coefficient for each ROI is shown separately. There was a significant positive correlation in the right fusiform cortex (*P < 0.005, Bonferroni corrected) but not in the other ROIs (n.s. P > 0.05, Bonferroni corrected).

## 4. Discussion

This study examined the hypothesis that attractiveness judgments on visual information degrades solely due to believing that the information is AI-synthesized regardless of its appearance. Participants underestimated face attractiveness when believing the faces to be synthetic compared to believing them to be real. The degree of this individually varying attractiveness underestimation was associated with instruction-dependent changes in fMRI signals in the right fusiform cortex. Thus, these results provide behavioral and neural evidence to support the initial hypothesis.

The present results demonstrated that the attractiveness of artificial objects for humans is reduced solely by the humans’ belief that the objects are artificial, suggesting that appearance is not the only factor influencing the attractiveness of artificial objects. By contrast, previous studies in AI/robotics have implicitly assumed that improving the attractiveness of AI/robots primarily based on their appearance (Abubshait & Wiese, 2017; DiSalvo et al., 2002; Gong, 2008; Kanda et al., 2008; Koda & Maes, 1996; McDonnell et al., 2012), relying on the idea from the uncanny valley hypothesis (Mori et al., 2012). Therefore, the present findings show the need to update the research assumption in AI/robotics, that improving appearance would suffice to improve the attractiveness of AI/robot-related visual information.

The analysis of brain responses revealed that, although multiple brain regions correlate to facial attractiveness (Fig. 3), only the right fusiform cortex changes its responsiveness to facial attractiveness together with the individual variation in attractiveness underestimation when participants believe that the faces they observe are synthetic (Fig. 4). The fusiform cortex contains a subdivision involved crucially in the visual processing of faces, namely the fusiform face area (Kanwisher et al., 1997), and contributes to the processing of facial attractiveness (Iaria et al., 2008). Meanwhile, neural signals in the fusiform cortex have been reported to distinguish artificial from living objects (Chaminade et al., 2010; Chao et al., 1999; Mahon et al., 2007; Noppeney et al., 2006). Moreover, a previous study on the neural mechanisms underlying the uncanny valley found that the human-likeness of artificial agents is signaled in the fusiform cortex (Rosenthal-von der Pütten et al., 2019). Therefore, the present results suggest that the neural signals of facial attractiveness and human-likeness interact and converge into the fusiform cortex. Particularly, I speculate that the reduced signals of facial human-likeness caused by believing faces to be synthetic produce the decrease in neural responsiveness to facial attractiveness in the fusiform cortex. This decreased responsiveness may weaken the perception of facial attractiveness, leading to attractiveness underestimation.

A recent study reported that humans judge synthetic faces as more trustworthy than real ones (Nightingale & Farid, 2022). Since facial trustworthiness is positively linked with facial attractiveness (Olivola & Todorov, 2017; Wilson & Eckel, 2006), this finding may appear inconsistent with the attractiveness underestimation observed in this study under the instruction of faces being synthetic. However, Nightingale and Farid compared the human trustworthiness of truly synthetic faces with that of truly real faces to uncover an intrinsic difference between synthetic and real faces. In contrast, this study aimed to elucidate how the human belief of faces being synthetic purely affects the attractiveness rating of the faces. Thus, as these two studies performed research from different perspectives, their findings are not necessarily inconsistent.

In this study, synthetic faces were used to explore the mechanism by which the attractiveness of that information is decreased due to the human belief that the information is AI-synthesized. However, faces are biologically salient objects, processed through a dedicated system in the human brain (Tsao & Livingstone, 2008). Therefore, whether similar behavioral and neural profiles can be observed with other types of visual information instead of faces remains unknown. Recent developments in generative AIs allow us to synthesize a variety of high-quality visual images other than faces (Ramesh et al., 2022; Rombach et al., 2022). Such visual images can also be examined through the attractiveness-rating task presented in this study (Fig. 1). Thus, it will be of interest to investigate the underestimation in attractiveness judgments using various types of visual information.

## 5. Conclusion

Our findings provide novel behavioral and neural evidence to support the hypothesis that the attractiveness judgments of humans on visual information degrades solely due to believing the visual information is AI-synthesized regardless of its appearance. A possible major factor for this attractiveness underestimation is thought to be the potential aversion of humans to AI. Thus, the present findings encourage future research in AI/robotics to explore how the potential human aversion to AI/robots can be dispelled to improve their attractiveness.

## Acknowledgements

I thank Ms. Hitomi Koyama for her experimental support. The work was supported by JSPS KAKENHI Grant-in-Aid for Scientific Research B (21H03535) and for Challenging Exploratory Research (22K19819), and JST PRESTO (JPMJPR20C6) to SN.

## Supplementary materials

**Fig. S1.**
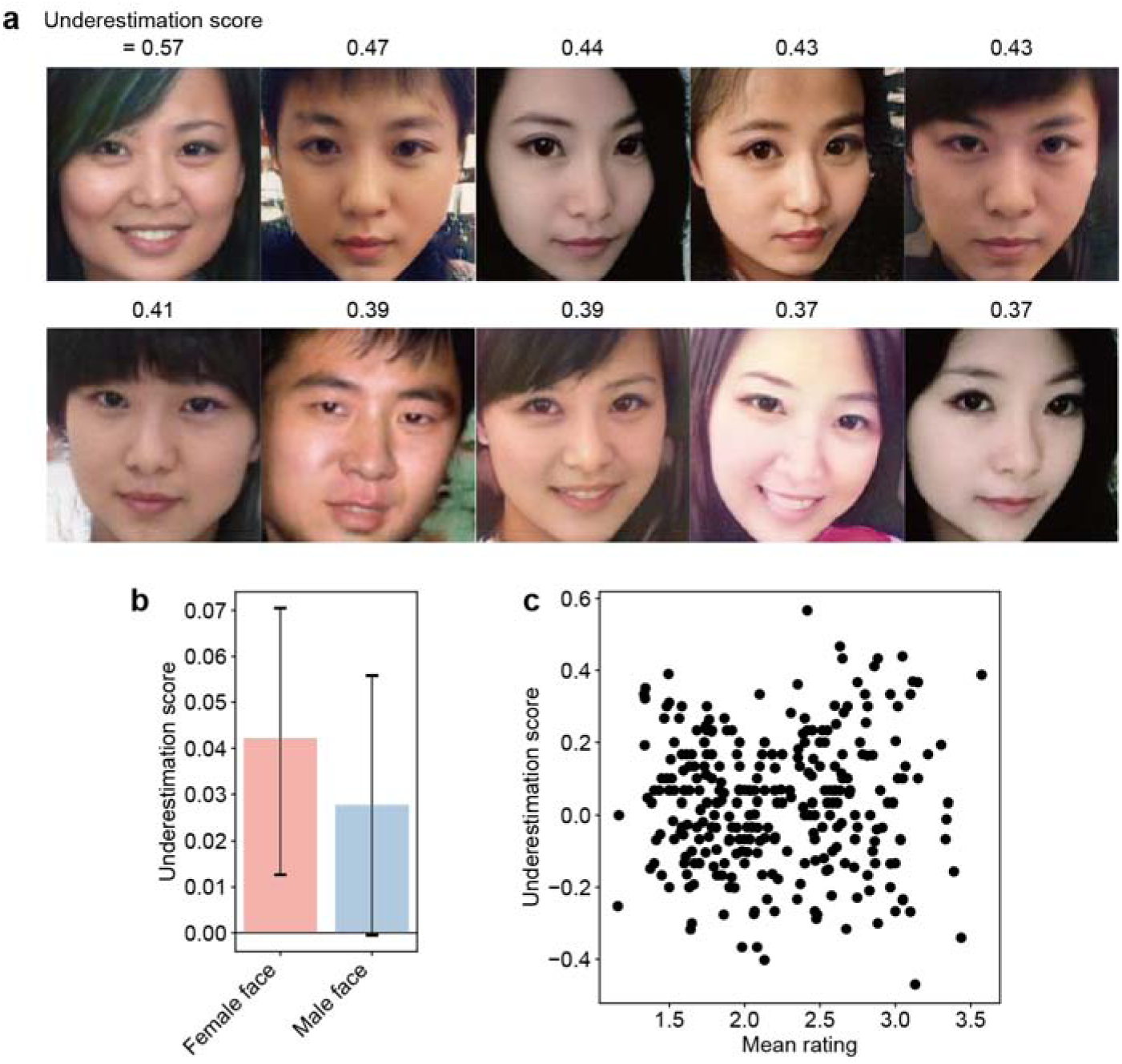
No clear tendency of facial attributes associated with underestimation scores. (a) Top 10 faces showing the highest underestimation scores. The value above each face indicates its underestimation score. The face images shown in this figure are AI-synthesized and not of real people. (b) Comparison of underestimation scores between faces of different sex (red, female; blue, male). Error bars indicate 95% CI. Sex differences in underestimation scores were not significant (two-sample t-test, p = 0.49). (c) Relationship between attractiveness ratings and underestimation scores. For each face (denoted by dots), the participant-average attractiveness rating (x-axis) and underestimation score (y-axis) are shown. There was no significant correlation between these two values (Pearson r = 0.021, p = 0.71).

**Fig. S2.**
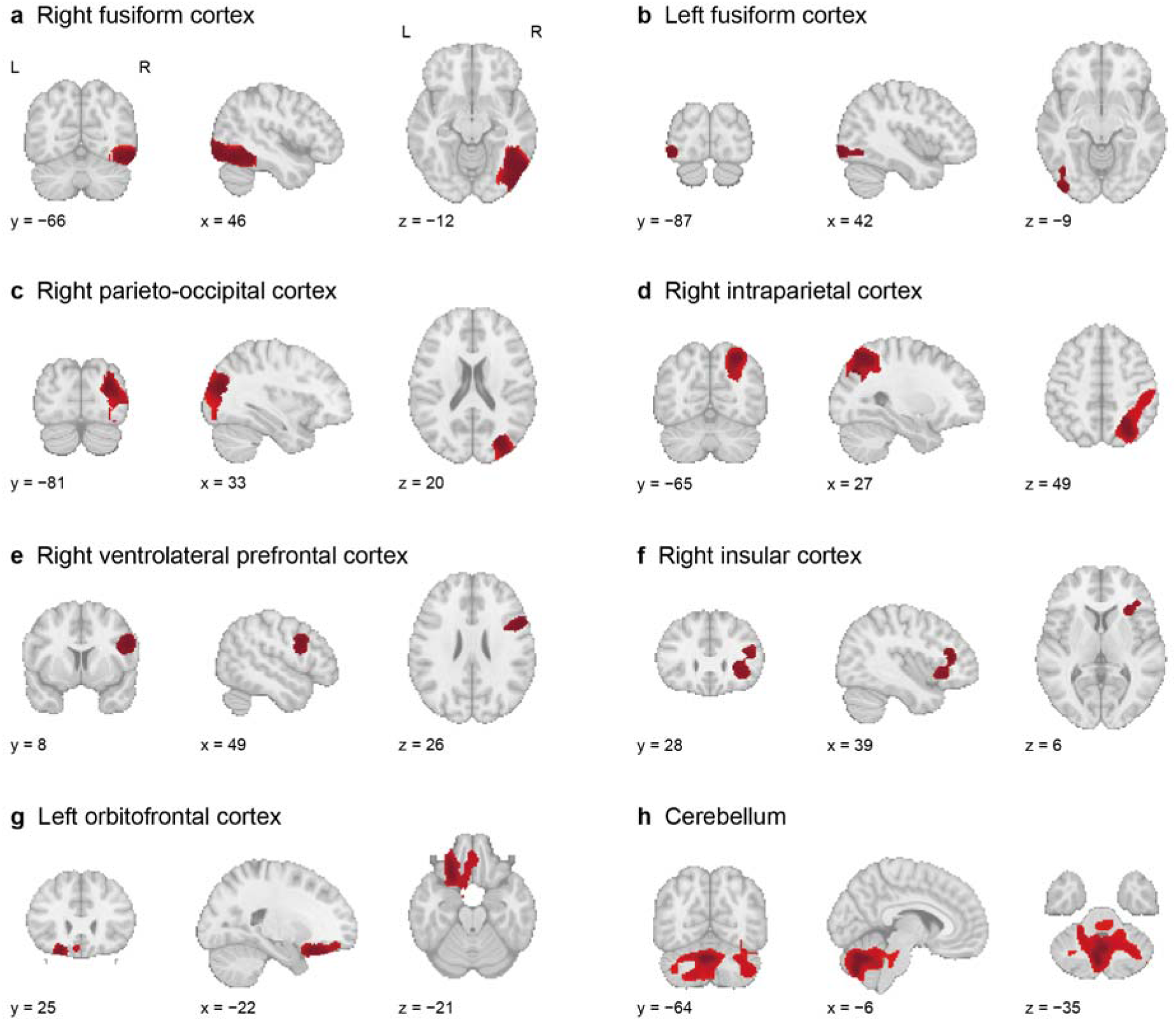
Eight brain regions used for the ROI analysis. Eight brain regions were identified as ROIs using second-level GLM analysis, which was performed to localize the voxels showing significant responsiveness to facial attractiveness. Each region consisted of connecting significant voxels forming a volume above a predefined threshold (400 mm^3^; for more details, see Materials and Methods). The eight regions, indicated by red-filled areas, were the bilateral fusiform cortex (a and b), the right parieto-occipital cortex (c), the right intraparietal cortex (d), the right ventrolateral prefrontal cortex (e), the right insular cortex (f), the left orbitofrontal cortex (g), and the cerebellum (h).

**Fig. S3.**
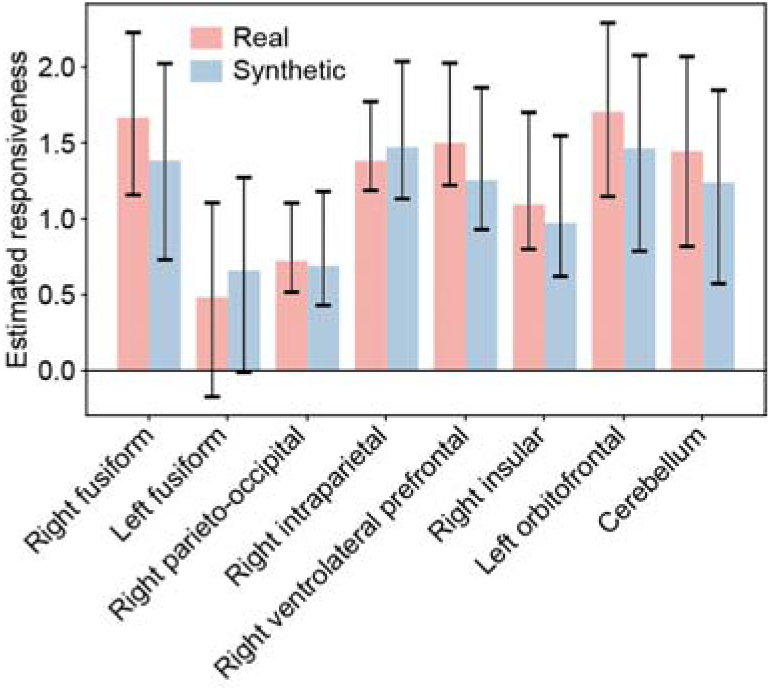
Condition-dependent responsiveness change in the eight ROIs. Estimated responsiveness to facial attractiveness were compared between synthetic- and real-face conditions at the group level. Bars represent group-average estimated responsiveness in each ROI under real- (red) and synthetic-face (blue) conditions. Error bars represent 95% CI. No ROI showed a significant response difference (paired t-test, P > 0.36, uncorrected).

